# Airway epithelial SARS-CoV-2 infectious and repair responses: relationships to age, sex, and post-COVID pulmonary syndromes

**DOI:** 10.1101/2025.07.17.663733

**Authors:** Hong Dang, Caitlin E. Edwards, Takafumi Kato, Boris Reidel, Rita M. Meganck, Charles R. Esther, Camille Ehre, M. Leslie Fulcher, Alexis B. Bailey, Michelle R. Cooley, Yu Mikami, Takanori Asakura, Padraig E. Hawkins, Minako Furusho, Jeffrey L. Myers, Kristine Konopka, Firoozeh V. Gerayeli, Hye Yun Park, Don D. Sin, Alessandra Livraghi-Butrico, Kenichi Okuda, Raymond J. Pickles, Sabra Klein, Scott H. Randell, Wanda K. O’Neal, Ralph S. Baric, Richard C. Boucher

## Abstract

The long-term pulmonary sequelae of SARS-CoV-2 respiratory infections reflect infection severity, innate and adaptive immunity, and respiratory epithelial repair. This study investigated the acute and reparative responses as a function of age and sex in primary human bronchial epithelial (HBE) cultures utilizing a 14-day SARS-CoV-2 infection protocol. SARS-CoV-2 infection peaked at 3 days post-infection (dpi) with an ∼ 2 log titer suppression at 14 dpi. SARS-CoV-2 infection induced interferon, interferon-induced gene, and cell damage responses. No age- or sex-dependent effects on SARS-CoV-2 infection were detected. Airway epithelia repaired to an abnormal mucus metaplastic/inflammatory state that reflected potentially beneficial and adverse consequences at 14 dpi. Repair processes were infection severity-dependent, not sex-dependent, and were more robust in young donor cultures. Analyses of long-COVID subjects with persistent pulmonary fibrosis or persistent bronchitic airway diseases exhibited expression of HBE 14 dpi failed repair gene signatures, including ISG gene signatures. Human airway epithelial repair post-SARS-CoV-2 is prolonged and incomplete *in vitro* over 14 days, and persistently abnormal repair may contribute to phenotypes of people with long-COVID pulmonary syndrome.

## Introduction

The respiratory outcomes following SARS-CoV-2 infections reflect multiple genetic and environmental variables (1–3). The severity of the lower airway infection, coupled with virus-modified epithelial repair programs and pre-existing epithelial health, are major variables predicting the severity of SARS-CoV-2 outcomes (4–8). Age and sex have also been related to SARS-CoV-2 disease severity (9–13). Children of both sexes are reported to exhibit less severe disease outcomes than adults (14, 15). Within the adult population, SARS-CoV-2 infection in older males generally is associated with more persistent disease (16). However, it has been difficult to attribute the relative risk of persistent disease in older males to differences in intrinsic epithelial infectivity/innate defense, age of the immune system, and/or comorbidities (17–20).

The first goal of this study was designed to identify potential contributions of human bronchial epithelia (HBE) to the acute and chronic (persistent COVID) outcomes of SARS-CoV-2 respiratory infections *in vitro* as a function of sex and age. Well-differentiated HBE cultures derived from sex-matched donors comprising a wide spectrum of age, including children aged 4 mo-8 yrs, young adults (20-30 yrs; referred to as mature), and the elderly (> 69 yrs), were studied over a prolonged post-SARS-CoV-2 inoculation (14 day interval) to capture potential age- and sex-dependent differences in: 1) the kinetics, magnitude and clearance of the initial infection; 2) the response/effects on the respiratory epithelium at peak viral infection; 3) how respiratory epithelia manifest SARS-CoV-2 infection post the peak infection period; and 4) how respiratory epithelia adapt and repair after peak epithelial infection/damage. A second goal of this study investigated whether *in vitro* d14-derived airway epithelial repair gene expression profiles persisted in specimens/data from subjects with prolonged (many months) failed epithelial repair manifested in fibrotic and “bronchitic” sequelae of SARS-CoV-2 infection.

## Results

### A. Global kinetics of virus infection

#### Kinetics of Viral Titers

Evidence of productive viral infection was present in 23 of the 24 experimental cultures of human bronchial epithelial (HBE) cells as assayed by live virus titers outlined in Protocol 1 (**Figure 1A and Supplemental Figure 1A;** note, the absence of infection of the one culture was deemed to reflect technical reasons, and it was removed from all analyses; see **Supplemental Table 1**). Measurable increases in virus titer were observed at 1 day post-infection (dpi), and titers peaked in 22/23 HBE donors at 3 dpi, the rate increasing with an average slope of 1.59 log titer/day (**Figure 1A and Supplemental Figure 2Ai**). Titers were reduced by 7 dpi and either leveled off or continued to decrease by 14 dpi, with 9 of 23 cultures exhibiting no measurable live virus titer at 14 dpi (**Figure 1A**). Consequently, throughout this report, cultures that exhibit a loss of live virus by 14 dpi are termed “suppressors” and those cultures with measurable titers still present at 14 dpi are termed “non-suppressors”. The slopes describing the decline of infection over the 3-14 dpi interval described a viral clearance rate ∼5 times slower than the rate of virus production between 1-3 dpi (average slope of -0.3 log titer/day; **Supplemental Figure 2Aii**). The rate of decrease was faster (by -0.23 log titer/day) in suppressors vs. non-suppressor HBE donors (**Supplemental Figure 2Aiii**). Collectively, the kinetics and magnitude of virus growth observed *in vitro* resembled those of previous *in vitro* studies (21, 22) and *in vivo* challenge studies of young healthy subjects (23).

**Figure 1.**
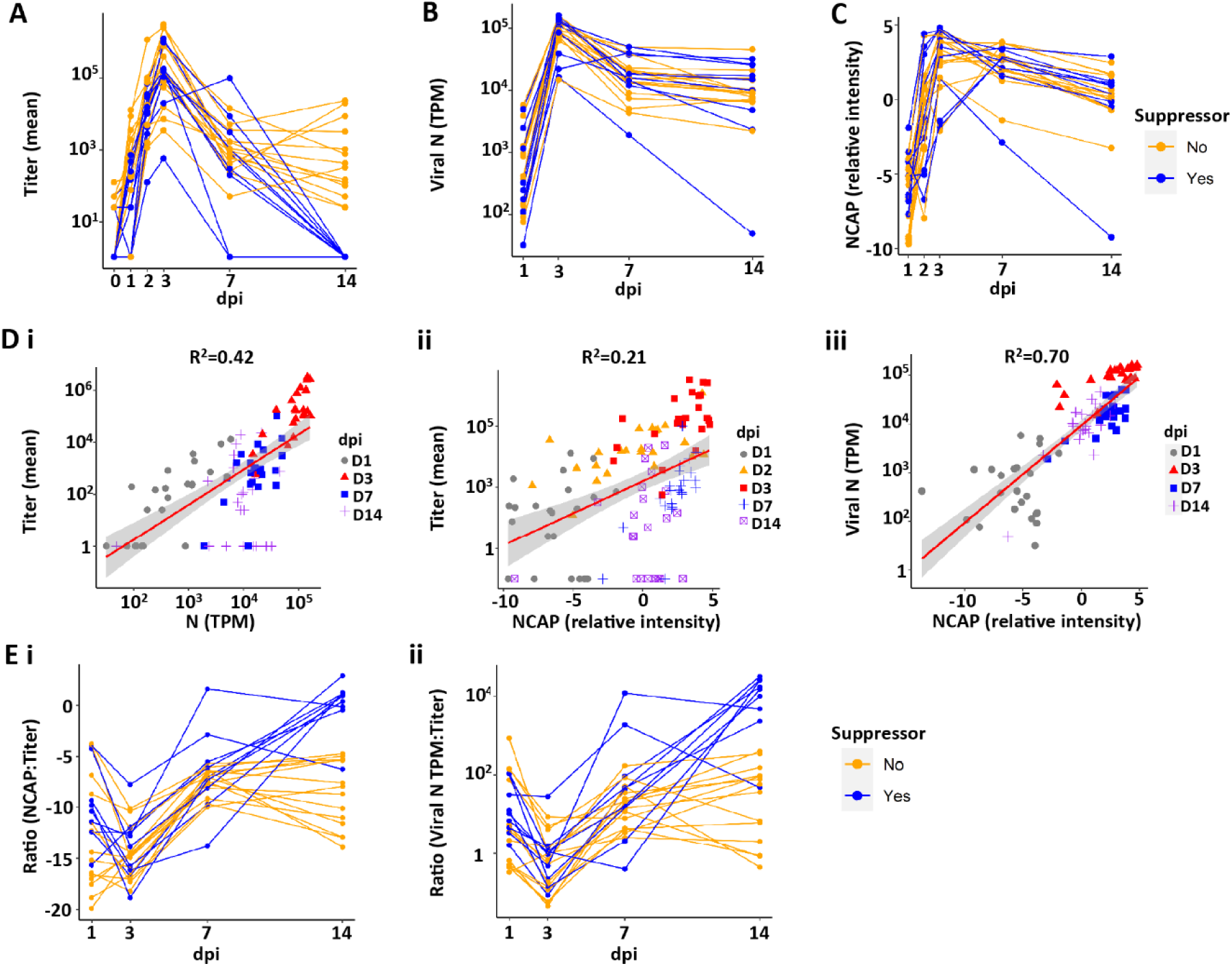
Indices of SARS-CoV-2 infection of all human bronchial epithelial (HBE) donor cultures with day post-infection (dpi). **(A)** SARS-CoV-2 titers over the 14 dpi interval, with each donor shown as connecting lines. HBE donor cultures that “suppressed” viral titers at d14 shown in blue. “Non-suppressors” are shown in orange. **(B)** Intracellular SARS-CoV-2 nucleoprotein (N) transcripts per million (TPM) by dpi for each donor culture. **(C)** Luminal extracellular viral NCAP protein by dpi for each donor. **(D)** Correlations between indices of SARS-CoV-2 infection. Shaded areas represent the 95% confidence interval. **(Di)** Correlation between viral titer and viral N TPM for each dpi. p < 0.0001. **(Dii)** Correlation between titer and nucleocapsid protein (NCAP) levels. p < 0.0001. Note that several 14 dpi samples are NCAP-positive and titer-negative. **(Diii)** Correlation between NCAP protein and SARS-CoV-2 viral N TPM for each dpi. p < 0.0001. Pearson correlation tests were used to calculate p-value and R^2^. **(E)** Relationships between indices of SARS-CoV-2 infection with dpi. **(Ei)** Ratio of total intracellular viral TPM vs. titer by dpi. **(Eii)** Ratio of NCAP protein vs. titer by dpi.

#### Kinetics of Total Intracellular Viral Transcripts

In addition to live virus titer determinations, intracellular viral RNA transcripts were measured at time of culture collection by bulk RNA sequencing (bulk RNAseq). The nucleocapsid (N) transcript was selected as an indicator of global SARS-CoV-2 viral mRNA and genome production (see Supplemental methods for explanation) (24). N transcripts, expressed as transcripts/million reads (TPM), were detectable by 1 dpi, increased to peak values at 3 dpi, and then subsequently declined through 14 dpi (**Figure 1B**). The rate of acquisition of viral TPM was similar to that measured by titer (slope 1.12 vs. 1.59, respectively; **Supplemental Figure 2, Bi vs. Ai**). In contrast to the loss of detectable live virus titer in suppressors, a complete loss of intracellular viral transcripts was not observed at 14 dpi in any HBE culture (compare **Figure 1, A and B, and Supplemental Figure 2, Aii and Bii, Aiii and Biii**). These data demonstrate the persistence of intracellular viral RNA in all cultures, despite the absence of measurable live virus in 39% of the cultures.

#### Kinetics of HBE Apical Surface Viral Proteome

Proteomics analyses identified SARS-CoV-2 viral proteins present in the apical washes sampled across all dpi (**Figure 1C and Supplemental Figure 2C**). Nucleocapsid (NCAP) was at the borderline of detection at 1 dpi, peaked at 3 dpi, and waned approximately 10-fold by 14 dpi (**Figure 1C and Supplemental Figure 2C, i and ii**).While we focused on NCAP protein as representative, similar patterns were observed for the spike protein, and, despite ∼10-fold less protein levels, for ORF8 (also called NS8) and ORF9B (**Supplemental Figure 2D**). As with intracellular viral TPM, extracellular viral protein was still detectable at 14 dpi in suppressor cultures (**Supplemental Figure 2Ciii**).

#### Correlations between Viral Titers, Intracellular TPM, and Apical Viral Proteins

Globally, viral titers were correlated with both SARS-CoV-2 RNA expression (using N mRNA as the representative transcript; **Figure 1Di**) and protein (using NCAP as the representative SARS-CoV-2 protein; **Figure 1Dii**). The correlation was strongest between SARS-CoV-2 RNA and NCAP protein (**Figure 1Diii**). The complete suppression of SARS-CoV-2 titers in cultures that continued to exhibit measurable SARS-CoV-2 gene expression and extracellular protein reduced the overall correlations with live virus titer. Comparisons of viral titer with SARS-CoV-2 TPM or extracellular viral protein with time revealed a reduction in both viral:TPM or viral:N protein from 1 dpi to 3 dpi, followed by a linear increase in ratios by 7 dpi, which leveled off in non-suppressors at 14 dpi (**Figure 1E**). Overall, these findings suggest two conclusions: 1) SARS-CoV-2 virus is maximally efficient at producing infectious virions in HBE cultures at 3 dpi, waning thereafter; and 2) a period of viral RNA persistence and protein shedding onto apical HBE surfaces in the absence of detectable infectious virions can occur late (*e.g.*, 14 dpi) in a SARS-CoV-2 infection, a finding of relevance to clinical antigen testing as a measure of infectious virus (23, 25).

#### SARS-CoV-2 Mutation Analyses

We tested the hypothesis that the mutation load in the SARS-CoV-2 genome would lead to the ability of the virus to persist in human airway epithelial cultures. Mutations acquired by the SARS-CoV-2 virus over 14 dpi were identified for each culture. Both coding and non-coding mutations were observed throughout viral RNA at 14 dpi **(Supplemental Figure 2Ei**). There were no differences in mutation load amongst cultures that suppressed or did not suppress viral titers (**Supplemental Figure 2Eii**). Thus, we conclude that HBE responses, not viral mutations, were the major drivers of viral kinetics in our system.

### B. Variables that affect HBE SARS-CoV-2 Infectivity

The basal state of airway epithelia is a major contributor to viral infectivity (26). Accordingly, we analyzed expression in 1 dpi mock exposed cultures for variables reportedly related to SARS-CoV-2 viral infectivity referenced to SARS-CoV-2 titers at 3 dpi. As expected, mRNA expression levels at 1 dpi in mock (basal) cultures for the proteins mediating SARS-CoV-2 binding/entry, *e.g.*, ACE2, TMPRSS2, and TMEM106B (27), were positively and significantly correlated with SARS-CoV-2 titers at 3 dpi (**Figure 2A-C**). Multiple studies have also focused on the relationships between ciliated cells and infection of airway epithelia by SARS-CoV-2 (22, 28–31). Two indices of HBE culture ciliation were utilized to examine relationships to infection: 1) The cilia-covered apical surface of the epithelium per basement membrane length was morphologically scored in mock cultures paired with infected cultures; and 2) a gene set based on cell ciliation was constructed and used to analyze mock 1 dpi cultures. The two metrics were correlated (r^2^=0.49; **Supplemental Figure 3A**), and positive but non-significant correlations were observed between both metrics and viral titer 3 dpi (**Figure 2D and Supplemental Figure 3B).** The relationships observed between viral titers with ACE2, TMPRSS2, and ciliation were generally similar when correlated to viral RNA (**Supplemental Figure 3C-E**).

**Figure 2.**
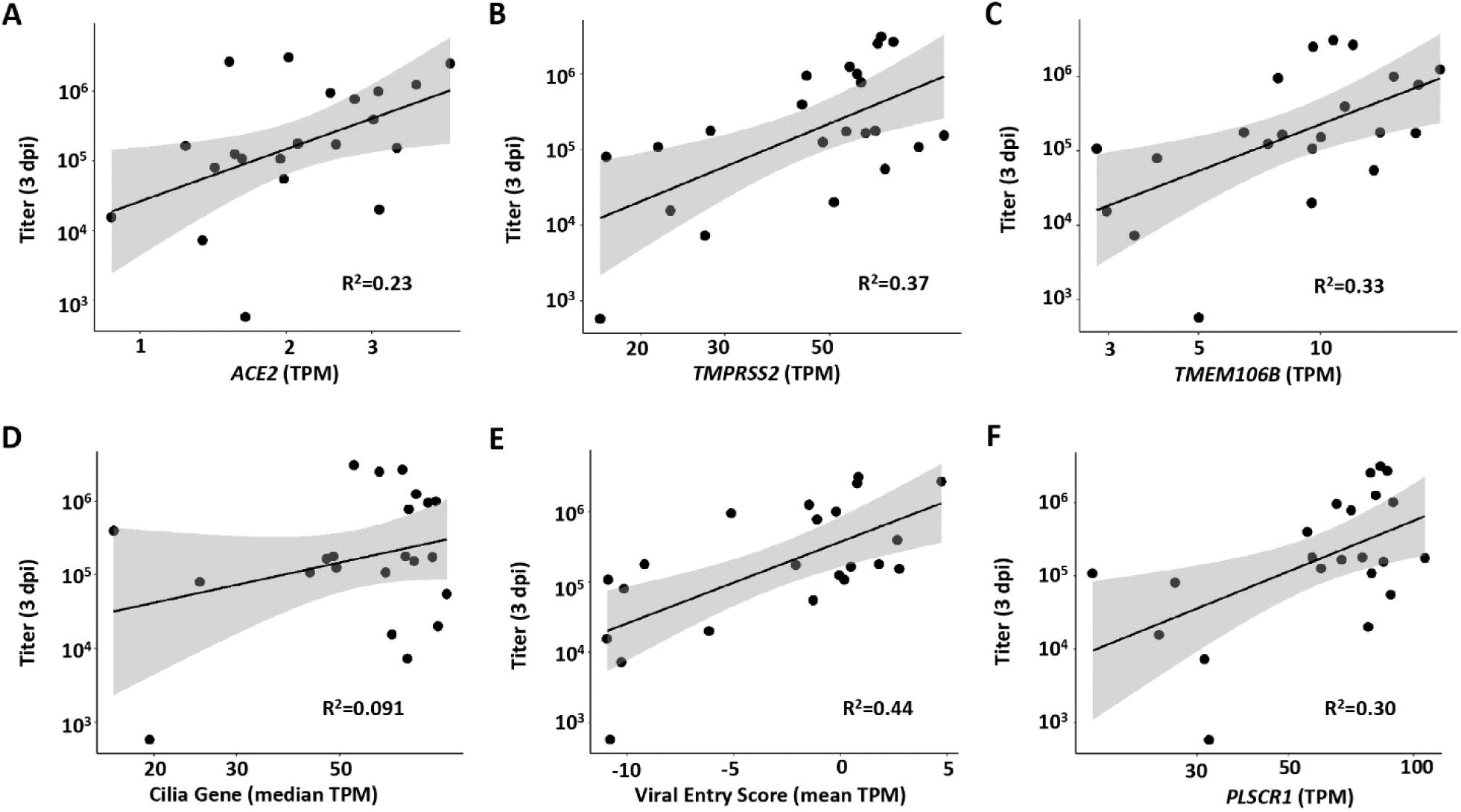
Host factor vs. peak viral titer relationships. Relationships between peak SARS-CoV-2 infection at 3 dpi and **(A)** long-form *ACE2* expression (p=0.24), **(B)** *TMPRSS2* expression (p=0.00249), **(C)** *TMEM106B* expression (p = 0.0049), **(D)** median cilia gene expression (p = 0.173), **(E)** HBE viral entry gene RNA score (p = 0.00071), and **(F)** *PLSCR1* expression (p = 0.0078). TPM=transcripts per million. All parameters listed on the X axes in A-F are measured from mock infected cultures at 1 dpi. Pearson correlation tests were used to calculate p-value and R^2^. Shaded areas represent the 95% confidence intervals.

We next evaluated whether baseline interferon expression, potentially reflecting a preexisting inflammatory state, would associate with viral titer at 3 dpi (26, 32–34). The baseline expression of a 13 gene index set of Type I interferon (IFN) stimulated genes previously reported, and with mean expression >50 TPM from mock infection samples, was utilized for this analysis (35). No relationships were detected between the basal expression of these genes and SARS-CoV-2 infection as indexed by viral titer or viral N gene TPM at 3 dpi (**Supplemental Figure 3F**). In addition, an index of genes reported to govern viral entry/infectivity was developed and tested for relationships to infectivity (36). A significant correlation was observed between basal expression of these genes and SARS-CoV-2 titer and viral N at 3 dpi (**Figure 2E and Supplemental Figure 3G)**. Curiously, many of these genes were also upregulated by virus in the bulk RNAseq data (**Supplemental Figure 3H**). Finally, PLSCR1 has been reported to inhibit SARS-CoV-2 entry/replication (37). In contrast to predictions, a positive correlation was observed between mock/basal PLSCR1 expression and SARS-CoV-2 titer at 3 dpi (**Figure 2F)**.

### C. IFN and ISG Response Kinetics

The interferon pathway is of critical importance in host responses to infection (26, 38–41). After SARS-CoV-2 infection, HBE cultures robustly increased expression of *IFNL1*, *IFNL2*, *IFNL3* and *IFNB1* (**Figure 3A**), with little expression or induction of *IFNL4, IFNA1, IFNG* or *IFNE* (**Supplemental Figure 4A**). The kinetics of the expressed IFN genes all followed a similar pattern, *i.e*., little induction was detected at 1 dpi, robust induction noted at 3 dpi, whereas induction waned rapidly by 7 and 14 dpi (**Figure 3A**).

**Figure 3.**
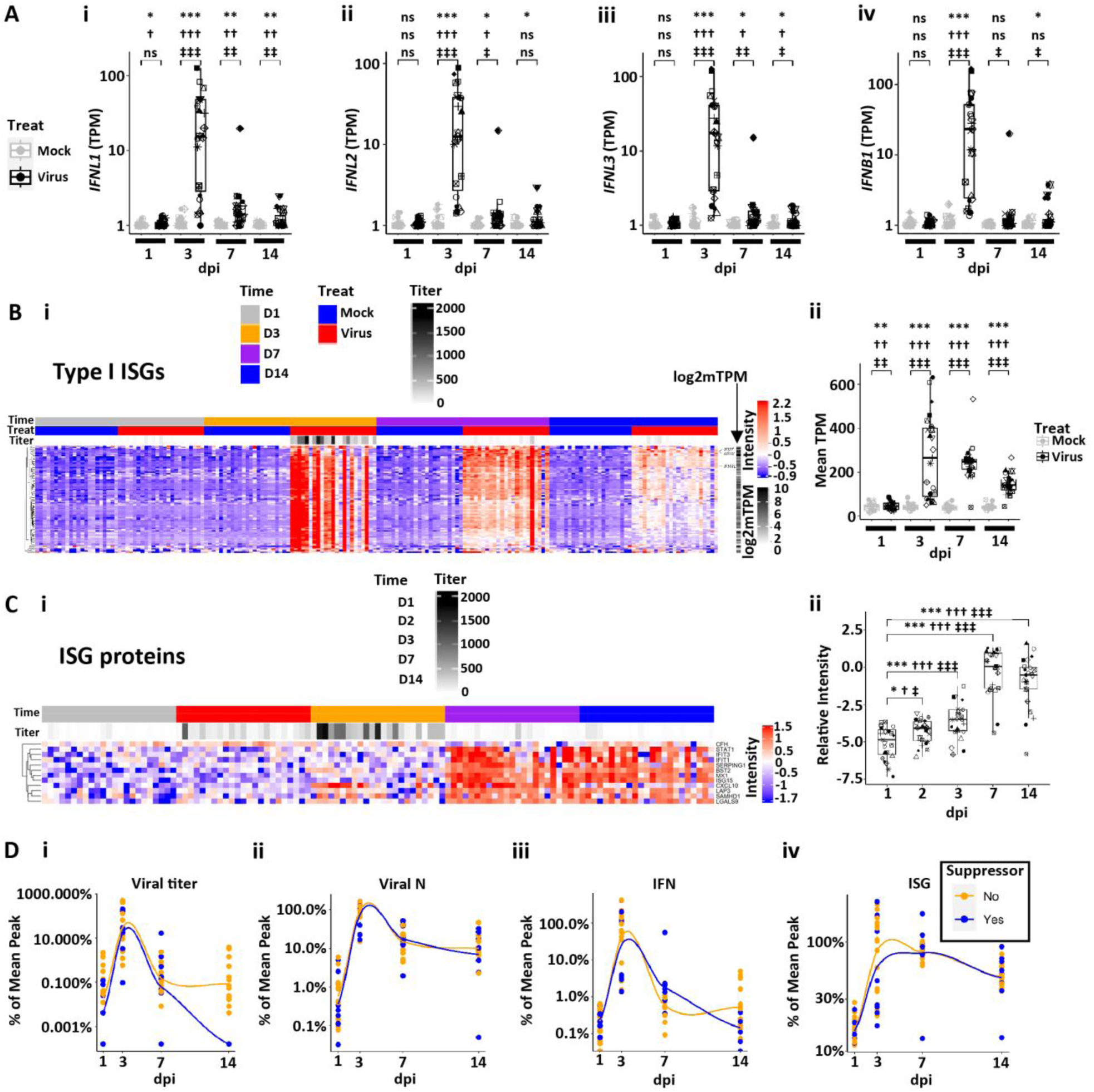
Interferon (IF) and interferon-stimulated gene (ISG) response kinetics to SARS-CoV-2 infection. **(A)** Box plots describing the expression values (in log10 scale plus offset of 1; TPM=transcripts per million) for HBE interferon responses over 14 dpi in mock and infected cultures for **(Ai)** *IFNL1,* **(Aii)** *IFNL2,* **(Aiii)** *IFNL3,* and (**Aiv**) *IFNB1***. (B)** HBE ISGs post-SARS-CoV-2 infection. **(Bi)** Heatmap describing pattern of ISG gene expression from 1-14 dpi. **(Bii)** Box plot describing level of mean ISG expression (TPM) for virus vs. mock from 1-14 dpi. **(C)** ISG proteins in luminal secretions from SARS-CoV-2-infected HBE cultures spanning 1-14 dpi. **(Ci)** Heatmap describing individual ISG proteins identified on HBE culture surfaces (normalized and imputed levels as described in methods). **(Cii)** Box plot describing mean level of secreted ISG protein for virus vs. mock from 1-14 dpi. **(D)** Relationships between viral infection, interferon (IFN), and ISG responses by dpi. Note, each parameter normalized to 100% mean peak response and depicted separately for HBE culture titer suppressors (blue) vs. non-suppressors (yellow). **(Di)** Mean percent of peak titers vs. dpi. **(Dii)** Mean percent nucleocapsid (N) transcript levels vs. dpi. **(Diii)** Mean percent IFN responses vs. dpi. **(Div)** Mean percent ISG responses vs. dpi. Genes associated with the heatmap are listed in **Supplemental File 1.** Statistics utilized linear mixed models using tiered statistical testing: a) Univariate gene expression (TPM) by treatment (using symbol *; top row), b) Linear models with biological variables (age group + sex + median ciliated genes) and treatment (using symbol †; middle row), c) Linear models with biological and QC variables (age group + sex + median ciliated genes + batch + PC1) and treatment (most conservative, using symbol ‡; bottom row). The number of symbols represents statistical significance: ns = not significant; one symbol = p<0.05; two symbols = p<0.01, three symbols = p<0.001; four symbols = p<0.0001.

Using a previously defined Type I INF-stimulated gene (ISG) set, a robust ISG response was observed that peaked at 3 dpi and waned thereafter (**Figure 3B**) (35). The ISGs remained elevated over baseline in most cultures at 14 dpi. Notably, a small number of ISGs exhibited a more persistent response, defined as greater fold changes over mock samples at 14 than 3 dpi.

These genes included *IFI27, BST2, IFI6*, and *IFI44L* (**Supplemental Figure 4B**). In contrast to RNA data, a smaller number of ISGs were detected by proteomics of luminal secretions (**Figure 3C**). ISG proteins markedly increased in secretions at 3 dpi above 1 dpi levels, including CXCL10, LAP3, and SAMHD1, whereas the majority of ISG proteins in luminal secretions peaked at 7-14 dpi. This difference may represent a lag between ISG gene induction and protein secretion and/or ISG release post-cell death.

To relate SARS-CoV-2 infection to IFN and ISG responses over the infection interval, we plotted SARS-CoV-2 infection, IFN responses, and RNA-measured ISG responses as a function of the percent peak mean response vs. time (**Figure 3D**). Compared to both viral titer and viral N TPM (**Figure 3D, i and ii**), the intracellular IFN RNA responses peaked at 3 dpi, waned over 7-14 dpi, and did not differ between viral titer suppressor vs. non-suppressor groups (**Figure 3Diii**). In comparison to IFN responses, the ISG responses exhibited similar kinetics at 3 dpi but were persistently expressed over the experimental interval (**Figure 3Div**), again with no significant differences between suppressor and non-suppressor cultures. These data are generally consistent with the associations predicted for a SARS-CoV-2-induced IFN and subsequent ISG response (41). Notably, the patterns of IFN and ISG responses were not directly related to the detection of live infectious virus (titer) at 14 dpi as revealed by suppressor vs. non-suppressor comparison. Thus, the persistent IFN and ISG responses could be driven by the continued presence of intracellular viral RNA transcripts and/or epigenetic changes that prevent a quick loss of IFN/ISG gene expression during a viral event.

### D. Global Transcriptomic Responses to SARS-CoV-2 Virus Infection 1-14 dpi

In addition to ISG responses, bulk RNAseq was used to catalogue other global epithelial responses to SARS-CoV-2 infection with time. Principal component analyses (PCA) and differential gene expression (DEG) analyses of the integrated HBE RNA expression dataset demonstrated that virus treatment was a strong overall driver of major differences in HBE cell gene expression post-infection at all timepoints measured (**Figure 4A-C**). SARS-CoV-2 infection induced significant up and down regulation of in gene expression, with different DEG noted across time post-infection, especially after 1 dpi (**Figure 4, C and D, and Supplemental Figure 5A**). A combined heatmap of differentially regulated genes comparing SARS-CoV-2-infected vs. mock HBE cultures across all times and treatment groups provides visualization of the major shifts driven by time (**Supplemental Figure 5B**).

**Figure 4.**
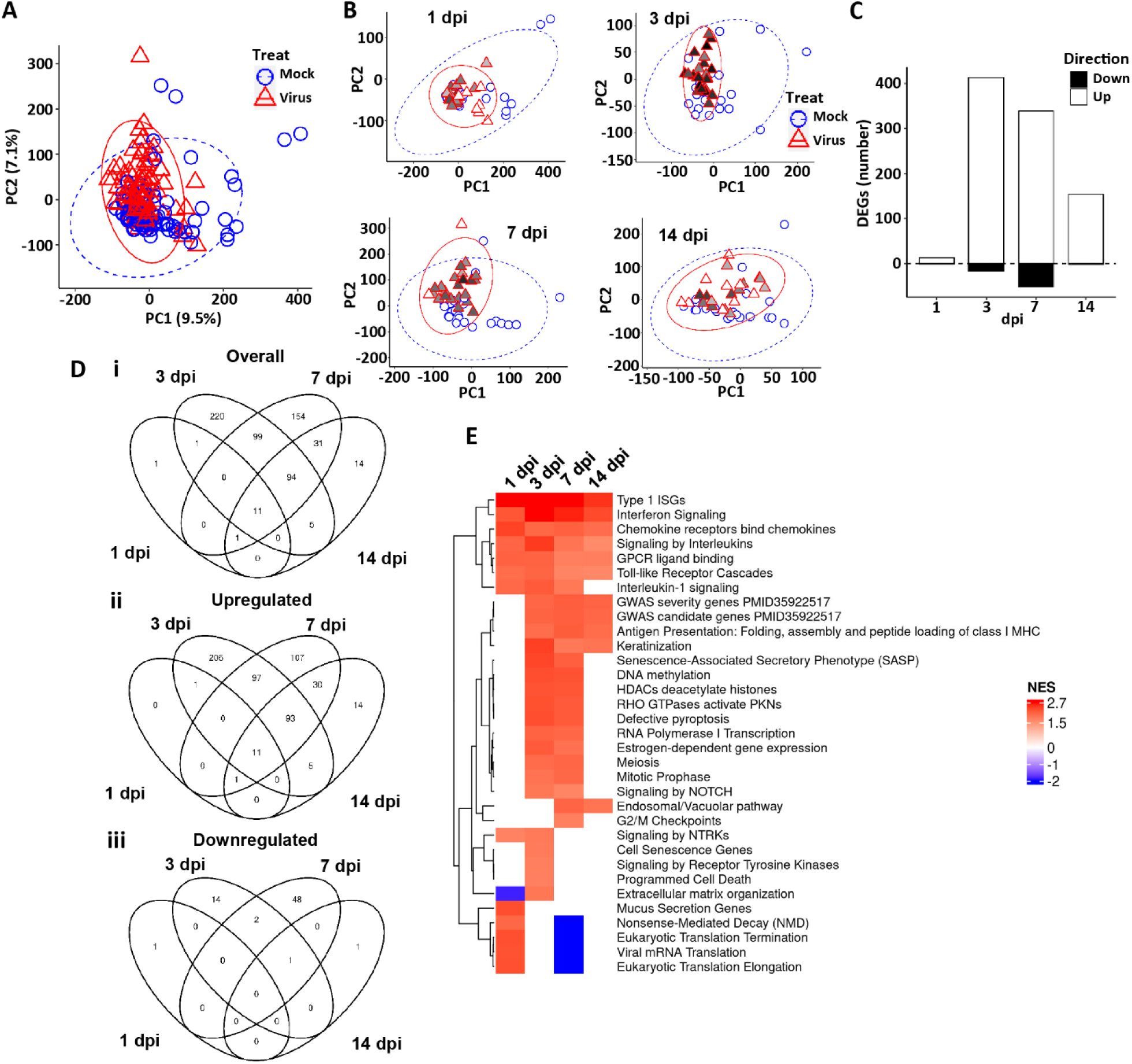
Global HBE responses to SARS-CoV-2 infection. **(A)** PCA plot representing the global bulk RNA transcriptome in mock and virus-infected cultures. All dpi are represented. **(B)** PCA plots representing global RNA transcriptome in mock and virus-infected cultures at the dpi indicated. In A and B, ellipses are drawn at 95% level assuming normal distributions of samples per group. PC=principal component. **(C)** Differentially expressed genes (DEGs) at the dpi indicated, showing number of regulated genes in both directions comparing virus-infected to mock treated cultures [(upregulated by SARS-CoV-2 in the upward bar (open); and downregulated in the downward bar (closed)]. DEGs represented are filtered at adjusted P-value of <0.05 and absolute log2 fold-change >1 with an average expression >0 (log2; or roughly TMP>1). **(D)** Venn diagrams depicting the overlap of DEGs between mock and SARS-CoV-2 infection across dpi groups. **(Di)** All DEGs; **(Dii)** DEGs upregulated by virus; **(Diii)** DEGs downregulated genes by virus. Filtering as in (C). **(E)** Heatmap of selected significant gene expression pathways identified as differentially expressed from the Reactome + custom pathway annotation set between virus infected vs. mock at indicated dpi. NES=normalized enrichment score. Red represents pathways upregulated by virus, blue represents pathways downregulated by virus.

As predicted from the large number of DEGs and the clustering data, pathway analyses identified highly significant pathway signatures driven by SARS-CoV-2 infection (**Supplemental Table 2**). A simplified heatmap of pathway responses demonstrated several key processes significantly altered at times from 1 to 14 dpi (**Figure 4E**). These pathways included interferon and cytokine signaling, mitotic signaling, NOTCH/Wnt signaling, and several stress pathways, including senescence and programmed cell death. Notably, no relationships were detected between IFs, ISGs, or genes associated with infectious viral production (*e.g.*, viperin, tetherin) and suppressor vs. non-suppressor status (data not shown). Down-regulated pathways at 7 dpi related to translation could reflect the activity of ISGs to control viral infection.

We also asked if genes identified in GWAS studies (42) were involved in epithelial responses to SARS-CoV-2 in HBE. Interestingly, many of the genes identified in GWAS studies associated with Covid-19 susceptibility and/or severity were expressed in airway epithelia, and as a group, they were responsive to virus infection after 3 dpi (**Supplemental Figure 5C**). These results suggest that the epithelial response, not just immune cell responses, to virus infection could play key roles in the presentation of Covid-19 in the clinic.

### E. Acute 1-3 dpi Cell Biologic Responses to Peak SARS-CoV-2 Infection of HBE Cultures

We next sought to characterize the effects of SARS-CoV-2 infection on HBEs at peak titers (3 dpi) using a combination of morphologic, cell biologic, proteomics, mass spectroscopy, and bulk RNAseq approaches. Previous studies have shown that the SARS-CoV-2 1-3 dpi interval is associated with a high degree of shedding of infected and apoptotic/necrotic dead ciliated cells (28). Cell death is also associated with release of cellular danger associated signals (DAMPs) (30). Consistent with these data, an increased incidence of cell death was observed at 3 dpi as indexed by the TUNEL assay (**Figure 5Ai and Supplemental Figure 6A**) and measures of *ZBP1* RNA (**Supplemental Figure 6B**) (43). Mass spectrometric analyses of luminal secretions revealed increased LDHB release over the 3-7 dpi interval (**Figure 5Aii**) and increased HMGB1 release that peaked at 3 dpi (44) (**Figure 5Aiii**).

**Figure 5.**
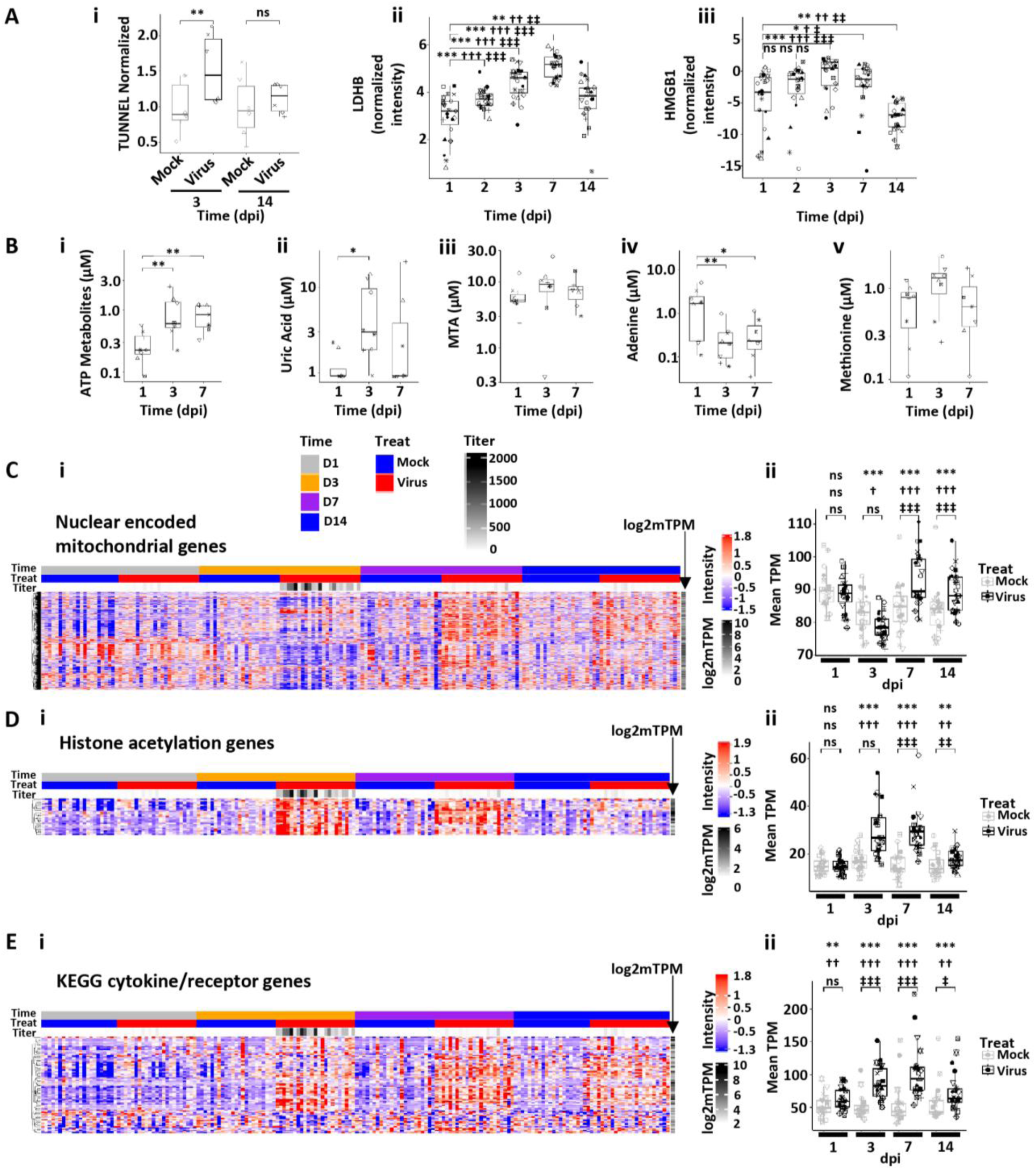
Human bronchial epithelial culture responses at peak SARS-CoV-2 infection. (A) Measures of cell death. **(Ai)** TUNEL assay quantification at 3 and 14 dpi in mock and infected cultures. ** = p < 0.01. **(Aii)** Box plots depicting LDHB release onto SARS-CoV-2-infected HBE culture surfaces spanning 1-14 dpi. **(Aiii)** Box plots depicting high mobility group box 1 (HMGB1) HBE apical surface protein levels spanning 1-14 dpi. **(B)** Measures of metabolites. Box plots depicting (**Bi**) total adenosine triphosphate (ATP) metabolites and (**Bii**) uric acid (UAC) concentrations, (**Biii**) MTA = methyl-thio-adenosine; (**Biv**) Ade = adenosine; and (**Bv**) Met = methionine as measured by mass spectroscopy in secretions at the indicated dpi. * = p < 0.05; **p<0.01. **(C-E)** Examples of pathway responses to peak SARS-CpV-2 infection. **(Ci)** Heatmap of nuclear-encoded mitochondrial genes post-SARS-CoV-2 HBE culture infection. **(Cii)** Box plots showing average gene expression per donor culture for mock and SARS-CoV-2-infected cultures over dpi. **(Di)** Heatmap describing histone acetylation gene expression over time post-SARS-CoV-2 infection. **(Dii)** Box plot describing average histone acetylation gene expression levels per HBE culture donor for mock and SARS-CoV-2-infected cultures with time. **(Ei)** Heatmap describing individual genes in the KEGG pathway for cytokines and receptors regulated by SARS-CoV-2 infection over dpi. **(Eii)** Box plots showing average gene expression per HBE donor culture for mock and SARS-CoV-2-infected cultures over dpi. Labels in Ci apply to all heatmaps. Statistics conducted and symbols as described in Figure 3 legend.

Airway epithelia also manifest a complex balance of small molecule-mediated mechanisms that can promote or dampen inflammatory responses to injury. We have previously reported that release of ATP as a DAMP is coincident with peak SARS-CoV-2 infections of HBE cells and serves as a second signal to promote inflammasome activation in locally resident airway macrophages (15). ATP is metabolized on airways surfaces by a series of extracellular metabolic enzymes that convert ATP to intermediate metabolites (AMP, adenosine, inosine, hypoxanthine, xanthine) with a final product of uric acid (UAC) (**Supplemental Figure 6C**). Expression of many genes within this pathway were altered in the 3-7 dpi interval (**Supplemental Figure 6Ci**), and mass spectrometric analysis of HBE apical secretions revealed increases in ATP metabolites and uric acid at day 3 dpi (**Figure 5Bi-ii and Supplemental Figure 6Cii**), consistent with acute ATP release serving as a pro-inflammatory/pro-inflammasome-activating moiety.

Another small molecule pathway known to be regulated in relationship to viral infection is the methionine salvage pathway, in which the anti-inflammatory molecule methylthioadenosine (MTA) is converted back to methionine (45). We observed an increase in apical concentrations of MTA with a corresponding decrease in adenine (a product of MTA hydrolysis) at day 3 dpi with no significant change in methionine levels (**Figure 5Biii-iv**) in parallel with decreased expression of genes involved in the methionine salvage pathway (**Supplemental Figure 6D**). These findings suggest the virus-induced downregulation of this salvage pathway with augmented MTA anti-inflammatory activity early in the post infection interval.

Bulk RNAseq analyses revealed additional global alterations in fundamental HBE cell biologic functions associated with peak SARS-CoV-2 infection. For example, a large fraction of the nuclear-encoded mitochondrial genes were suppressed at 3 dpi but were overexpressed later (**Figure 5C**). A strong upregulation at 3 dpi of DNA methylation/chromatin remodeling signatures, *e.g*., histone acetylation, was also observed which was maintained but declined over 14 dpi (**Figure 5D**).

Inflammatory/cytokine genes included in the “cytokine-cytokine receptor interaction” KEGG pathway (hsa04060) were also globally elevated by virus at 3 dpi and after (**Figure 5E**). However, the genes in this pathway appeared to cluster in two broad categories, one that peaked at 7 dpi and remained elevated and a second group that that were non-responsive to virus infection.

### F. Adaptive, Maladaptive, and Reparative 3-14 dpi HBE Responses Post-Peak Viral SARS-CoV-2 Infection

In many airway disease states following injury, *e.g*., virus infection, the epithelium remodels from the ciliated cell-dominated state to one characterized by mucus cell metaplasia (46). Accordingly, we characterized the cellular morphologies of HBE cultures over the post-SARS-CoV-2 peak infectious interval to test for repair/metaplastic processes post-SARS-CoV-2 infection. While ciliated cell loss was not evident at 3 dpi, an ∼ 50% reduction of ciliated cell numbers in infected cultures was observed at 14 dpi compared to mock (**Figure 6Ai**). Further, the ratio of ciliated cell gene expression to ciliated cell numbers differed between infected and mock cultures, with lower numbers of ciliated cells present for equivalent ciliated cell gene expression in the infected group (**Figure 6Aii**). A comprehensive heatmap of ciliated cell gene expression shows high variability, but a general reduction in mature multi-ciliated cell gene expression after virus infection was observed at 3 and 7 dpi (**Supplemental Figure 7A**). Newly differentiating ciliated cells transition from suprabasal/secretory cells through a deuterosomal cell state (47). Interestingly, despite an overall decrease in ciliated cells and ciliated cell gene expression, HBE cultures post-virus exhibited an increase in deuterosomal cell gene expression, especially at 7 dpi (**Figure 6Aiii and Supplemental Figure 7B**). These data suggest that there is a delay/failure of the deuterosomal cell population to regenerate a mature multiciliated epithelium.

**Figure 6.**
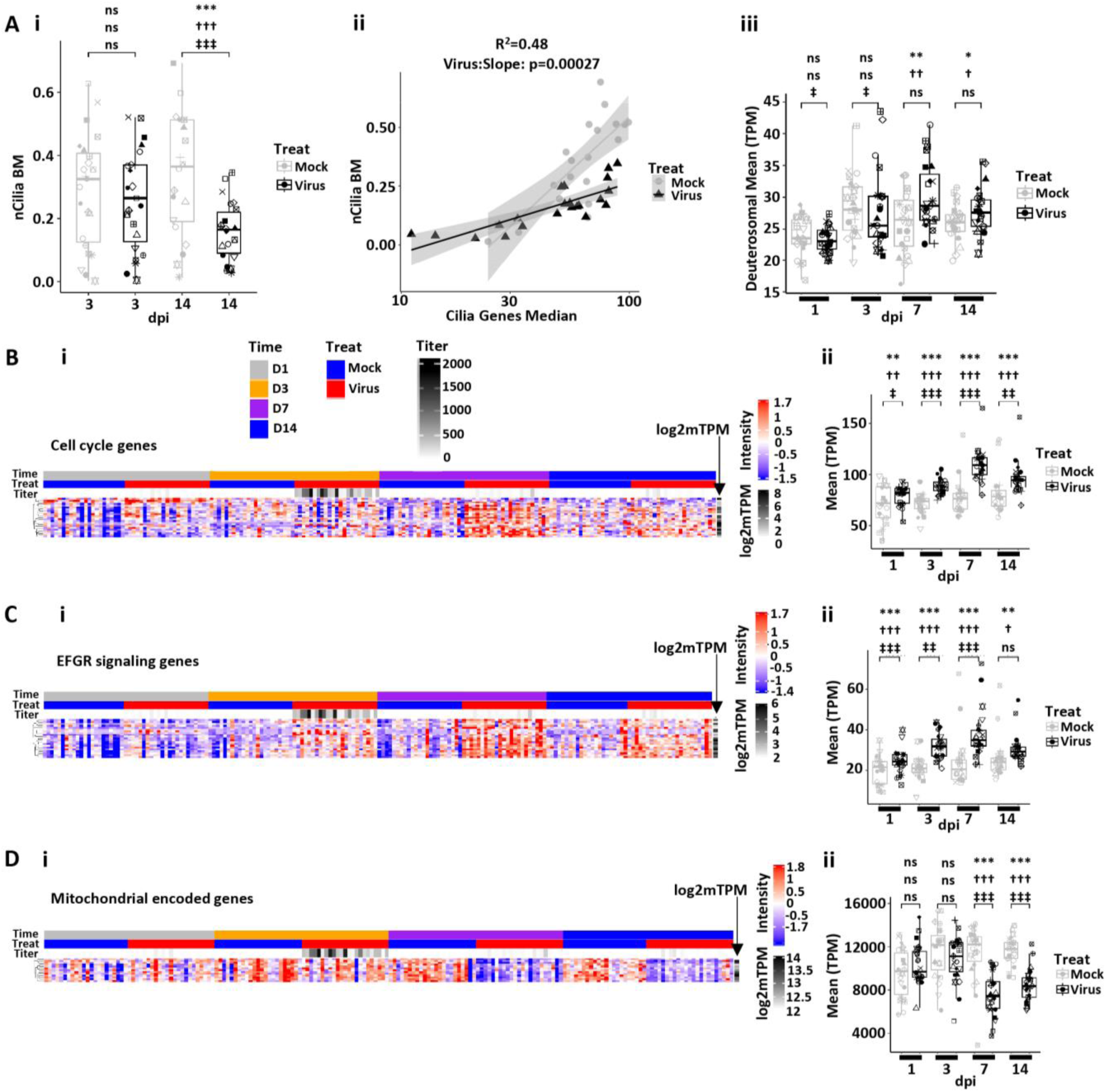
SARS-CoV-2-induced remodeling, senescence/inflammation, and repair in human bronchial epithelial cell cultures. **(A)** Ciliated cell quantification with infection. **(Ai)** Morphometric measurement of ciliation [cilia-covered apical surface/basement membrane (BM); measurement designated nCilia BM on axis label] for mock and SARS-CoV-2-infected HBE cultures at 3 and 14 dpi. **(Aii)** nCilia BM vs. median multi-ciliated cell gene expression pathway at 14 dpi. Mock HBE slope = 0.286 (SE 0.047), SARS-CoV-2-infected slope = 0.086 (SE 0.011) (delta = 0.176, SE = 0.043, p < 0.00027). Shaded areas represent the 95% confidence intervals. **(Aiii)** Mean deuterosomal cell gene expression for mock and SARS-CoV-2-infected cultures over dpi. **(B)** Cell cycling. **(Bi)** Heatmap depicting cell cycle gene expression over time post-infection. **(Bii)** Box plot depicting mean cell cycle gene expression (transcripts per million = TPM) over time post-infection. **(C)** EGFR signaling. **(Ci)** Heatmap depicting EGFR signaling genes as a function of time post-infection. **(Cii)** Box plots depicting mean EGFR signaling pathway gene expression over time post-infection. **(D)** Mitochondrial encoded genes. **(Di)** Heatmap depicting mitochondrial encoded genes over time post-infection. **(Dii)** Box plot depicting mean mitochondrial gene expression over time post-infection. Statistics conducted and symbols as described in Figure 3 legend. Headers in Bi apply to all heatmaps.

Consistent with our previous report, HBE cultures demonstrated an increase in AB-PAS-positive cells by 14 dpi, consistent with mucus cell metaplasia, in SARS-CoV-2-infected cultures (**Supplemental Figure 7C** (46). Mucus cell metaplasia can reflect metaplasia of pre-existing secretory cells into a mucus (“goblet”) cell phenotype and/or commitment of regenerative cells, *e.g*., basal cells/suprabasal cells, into this cellular phenotype. Pertinent to these possibilities, analyses of bulk RNAseq data revealed 1) an increase in basal cell and cell cycling genes (*e.g*., TP53 cell cycle pathway) in the interval post-peak infection, consistent with the notion that at least part of the mucus cell metaplasia represented a cell proliferative/replacement response to viral injury (**Figure 6B**); and 2) strong upregulation of EGFR signaling pathways that are likely to mediate a component of the mucus metaplastic response to SARS-CoV-2 (46, 48, 49) (**Figure 6C)**.

The post-infection 7-14 dpi HBE epithelium also exhibited other indices of chronic injury/damage. For example, upregulation of genes associated with cellular senescence at 3, 7 and 14 dpi was observed (**Supplemental Figure 7D**). Epithelial cells in the senescent state may contribute to the pro-inflammatory pathways that persisted over the 14-d interval post-infection (**Figure 5E**) (50). Adrenomedullin, which recruits ILC2s into airway walls and promotes Type 2 responses, was also upregulated in the post 3 dpi state (**Supplemental Figure 7E**) (51). In parallel, there was a reduction in mitochondrial-encoded gene expression over the 7-14 dpi interval, supporting the concept that virus infections lead to defects in cellular bioenergetics (**Figure 6D**). The downregulation of mitochondrial-encoded genes contrasted to increased expression of nuclear-encoded mitochondrial genes shown in **Figure 5C**. Thus, the post-SARS-CoV-2-infected epithelium exhibits a complex spectrum of repair, adaptation, and dysfunction.

### G. Host Defense Responses to SARS-CoV-2-Infected Airway Epithelia

In the context of injury, airway epithelia need to maintain elements of host defense. A major host airway defense mechanism is the mucociliary transport system (52). This system is in part dependent on ciliated cell numbers, which are reduced (see **Figure 6A**), predicting a degradation of the efficiency of this system (53). However, mucociliary transport is also governed by the mass of mucins on airway surfaces and the hydration of secreted mucins mediated by airway epithelial ion transport systems (52, 54). An increase in the dominant host defense airway epithelial secreted mucin, MUC5B, was observed by RNA, proteomics, and histologic measurements, particularly at the 14 dpi interval post SARS-CoV-2 infection (**Figure 7A**). *MUC5AC* was expressed basally at lower levels, its expression increased transiently with infection, but the absolute RNA levels remained low compared to MUC5B (**Supplemental Figure 8A**). Note that overall MUC5B RNA and protein levels, but not MUC5AC, correlated well (**Supplemental Figure 8B**).

**Figure 7.**
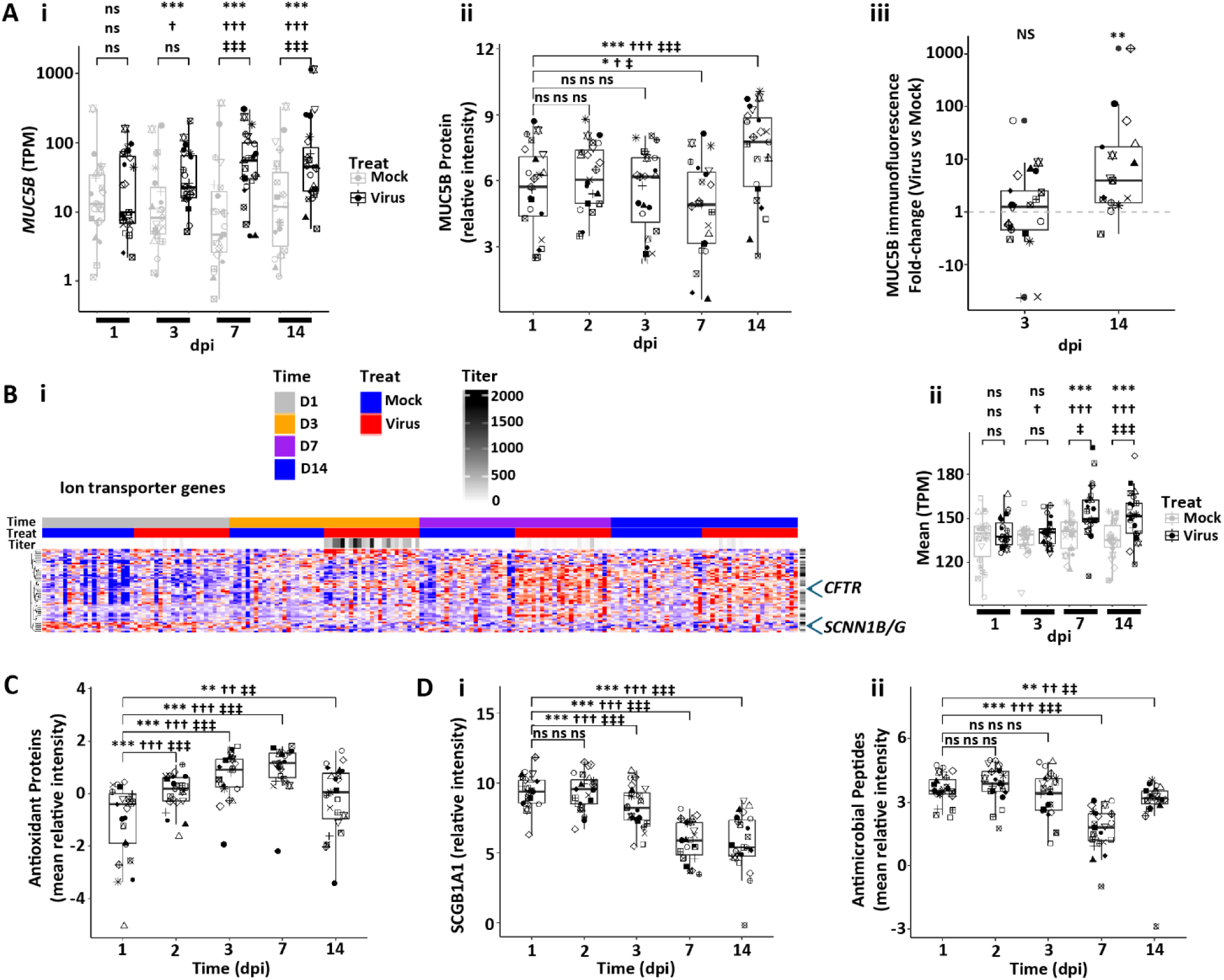
Human bronchial epithelial host mucosal defense responses to SARS-CoV-2 infection. **(A)** Secretory mucin expression. **(Ai)** Box plots depicting *MUC5B* RNA expression in mock vs. SARS-CoV-2-infected HBE cultures over dpi. **(Aii)** MUC5B protein secreted onto apical surfaces of mock vs. SARS-CoV-2-infected HBE cultures with time. **(Aiii)** Immunofluorescent MUC5B levels in SARS-CoV-2 ratioed to mock HBE culture levels at 3 and 14 dpi. **(B)** Ion transport genes. **(Bi)** Heatmap depicting pattern of ion transport genes in mock vs. SARS-CoV-2-infected HBE cultures with time post-infection. **(Bii)** Box plot depicting mean ion transport gene expression over time. **(C)** Box plot depicting mean antioxidant proteins collected from culture apical surfaces over time post-infection. **(D)** Antimicrobial defense molecules. (**Di**) Box plot depicting SCGB1A1 protein levels with time post infection. **(Dii)** Box plots depicting mean Reactome antimicrobial proteins with time post-infection. Statistics conducted and symbols as described in Figure 3 legend.

In parallel with increased mucin secretion, there was a significant upregulation at 7-14 dpi of a broad class of ion transport genes that mediate fluid secretion (**Figure 7B**). A central pathway mediating airway surface hydration includes the CFTR Cl^-^ channel and regulators of CFTR Cl^-^ channel fluid secretory activity (55). Both CFTR gene expression and multiple regulatory components of CFTR activity were upregulated. In contrast, key genes involved in fluid absorption, *e.g.*, *SCNN1β* and *SCNN1γ*, were not upregulated and clustered separately in the heatmap. The mucin and ion transport gene expression signatures suggest that airway surfaces post-infection upregulate pathways to maintain/restore mucociliary transport by secretion of a well-hydrated mucus to offset loss of ciliation and “flush” viruses/shed cells from airway surfaces.

Finally, airway epithelia can respond to infection with increased synthesis and secretion of host defense proteins and small molecules. While annotated antioxidant genes were variable in their response post-infection (**Supplemental Figure 8C**), secreted antioxidant proteins were routinely increased post-infection (**Figure 7C and Supplemental Figure 8D**). In contrast, proteomics revealed a striking reduction in secretion of other host defense proteins, *e.g.*, SCGB1A1 (**Figure 7Di and Supplemental Figure 8E**), and the pathway for antimicrobial peptides (which includes lysozyme, lactoferrin, SLPI, S100A8/9, and BPIFA/B1) was amongst the most downregulated proteomic pathway signatures across the entire study (**Figure 7Dii and Supplemental Figure 8F)** (56). Notably, no relationships were detected between the suppressor/non-suppressor state at the RNA or protein level for secreted proteins, with reported antiviral effects, including lactotransferrin, BPIFA1, BPIFB1, SLIPI, or MUC5B/MUC5AC mucins (data not shown). At the RNA-only level, no relationships were observed between the suppressor vs. non-suppressor state and beta, defensins, nitric oxide synthesis 1, 2, or 3, and cathelicidin (data not shown).

### H. Comparison of Infection of HBE Cultures as a Function of the Sex of the Donor

Note, because the sexes were not perfectly matched across batches in the 14 day protocol (**Supplemental Figure 1A**), a batch correction was necessary, which confounded the sex analyses (see **Supplemental Methods**). However, because, as described below, female sex was found to produce higher viral titers in the 14 day protocol which could have dominated over technical batch variables, the Protocol 1, 14 day studies were supplemented by data from a replicate single-batch 3 day study (Protocol 2) focused on the effects of sex on infectivity from 0-3 dpi (**Supplemental Figure 1B**). Each protocol is discussed separately below.

In the 14 day protocol (Protocol 1), female donor HBE cultures exhibited a more rapid rise in SARS-CoV-2 infection and higher peak levels of SARS-CoV-2 infection than cultures from male donors as indexed by viral titers and secreted NCAP protein (**Figure 8, Ai and ii, Supplemental Figure 9A**). In contrast, the mean rates of suppression of viral titer, N TPM, and extracellular viral proteins were not different between females and males (**Figure 8A, iii and iv, and Supplemental Figure 9A and B**). Differences in cilia, ACE2, TMPRSS2 and TMEM106B were not evident between the sexes that would explain the increase viral titer in females at 3 dpi (**Supplemental Figure 9, C and D**). No differences were observed in SARS-CoV-2 mutation load as a function of sex (**Supplemental Figure 8E**).

**Figure 8.**
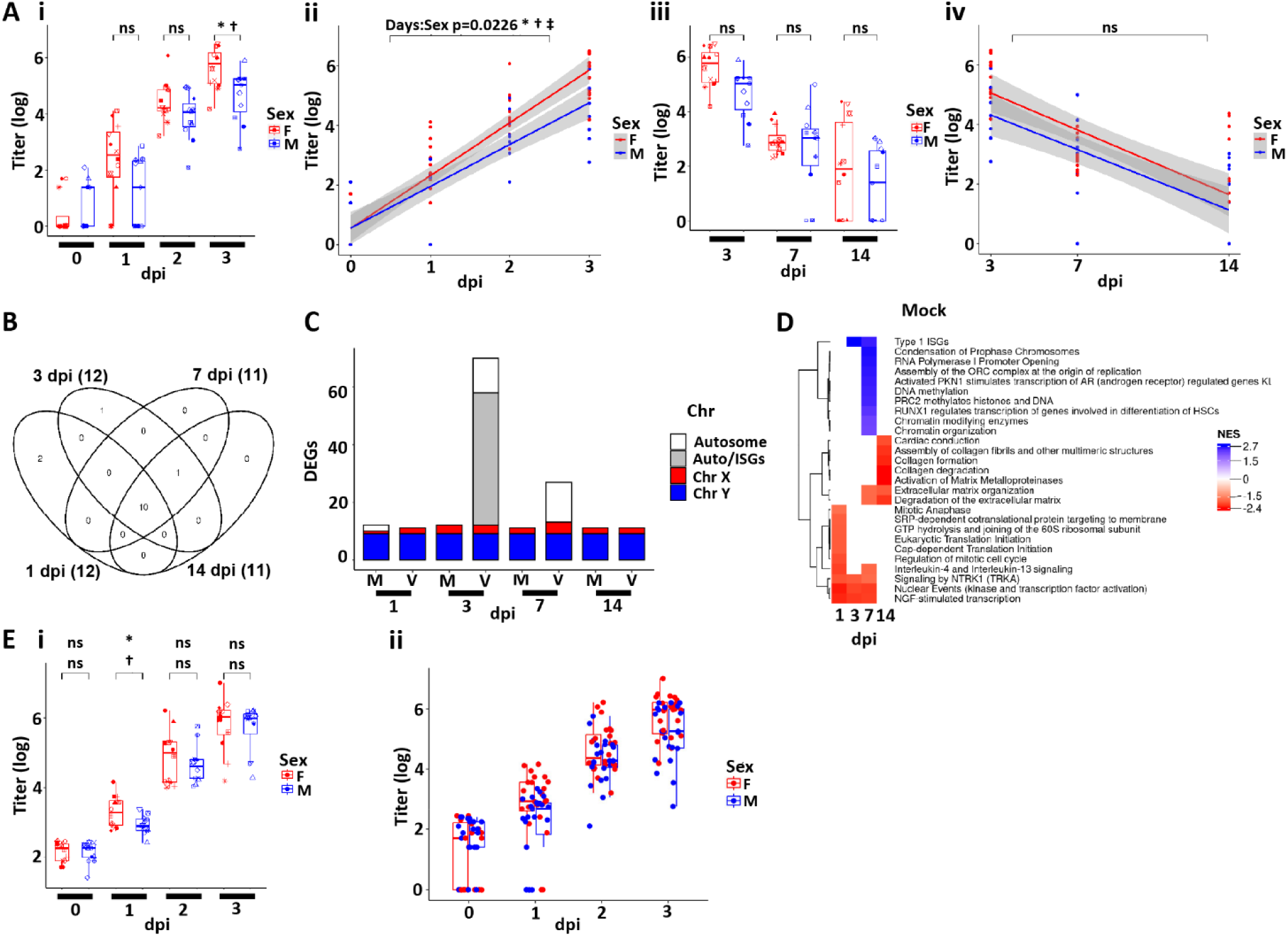
SARS-CoV-2 infectivity and HBE culture host responses as a function of sex. **(A)** Box plots depicting mean titers in SARS-CoV-2 infected HBE cultures from 0-3 dpi colored by sex. **(Aii)** Slopes of mean titers in each sex for dpi 0-3 (slope = 1.77 for females and 1.40 for males). The p-value of the difference between the slopes (-0.37) is 0.0226. The shaded areas represent the 95% confidence intervals. **(Aiii/iv)** As for (Ai-iii) expect representing titer data for dpi 3, 7, and 14. **(B)** Venn diagram representing differentially expressed genes (DEGs) between sexes from analyses of mock HBE cultures at the dpi indicated. Numbers in parentheses represent the total number of DEGs in each group. DEGs represented are filtered at adjusted P-value of <0.05 and absolute log2 fold-change >1 with an average expression >0 (log2; or roughly TMP>1). **(C)** Schematic representation of DEGs between the two sexes at the times and treatments indicated (m=mock; v=virus). Coloring represents the location of the DEGs on the chromosomes where Auto=all autosomes. DEGs represented are filtered as for (B). **(D)** Simplified heatmap of differentially expressed pathways from Reactome + custom annotation set for female vs. male in mock infected HBE cultures. Red pathways are upregulated in females compared to males; blue pathways are upregulated in males compared to females. NES=normalized enrichment score. **(E)** Replication cohort (Protocol 2). **(Ei)** Box plots depicting mean titers in SARS-CoV-2 infected HBE cultures from 0-3 dpi colored by sex Protocol 2. **(Eii)** Combined titer data for Protocols 1-2. No significant difference between sexes were found for any timepoint measured.

To test for sex-dependent differences in host epithelial genes that might relate to sex-dependent differential SARS-CoV-2 infectivity, global transcriptomic differences between male and female mock cultures were evaluated to characterize the state of gene expression prior to infection. Notably, very few differentially expressed genes were observed between male vs. female cultures over the 1-14 dpi interval in the mock infected cultures (**Figure 8B**). Nearly all the differentially expressed genes in the absence of infection (mock) between female and male cultures reflected the sex chromosome genes (**Figure 8C**), and no pathways were consistently different in male vs. female mock cultures across the four timepoints measured (**Figure 8D**). The major difference between the sexes after virus infection were noticed as an increase in the number of genes upregulated in females at 3 dpi (**Figure 8C**). Further evaluation of those genes identified them as ISGs, which correlated with the increase in viral titer in females at 3 dpi. These ISGs were not differentially regulated between male and female 14 dpi when viral titers were not different. We conclude that in this study, male and female differences in the airway epithelial transcriptome were extremely rare, dominated by X-Y chromosome-specific genes, with no one expression profile predicting differences in infectivity.

Given the importance of possible sex differences in SARS-CoV-2 infectivity observed in the first protocol, the acquisition phase of infection in HBE cultures was repeated for same donors but infected as a single batch (**Supplemental Figure 1B**, Protocol 2). In the replication study, while similar trends for females to show early increases in infectivity, at 1 dpi, no differences were observed at 2 and 3 dpi (**Figure 8Ei**). Importantly, the combined data from the two studies revealed no significant difference in infectivity between the sexes (**Figure 8Eii**). Thus, we conclude there was little significant difference in sex-related SARS-CoV-2 infection of human airway epithelia.

### I. SARS-CoV-2 Infection and repair of HBE cultures as a function of donor age

The 14 day two-batch protocol (Protocol 1) was matched for age. There were no differences in the rate of acquisition of SARS-CoV-2 infection over the 0-3 dpi interval as a function of age as reflected in titer, N TPM, or extracellular N (**Figure 9, Ai and E, and Supplemental Figure 10A**). Further, there were no consistent differences in resolution over that 3-14 dpi interval as a function of donor age despite a tendency for young cultures to be more likely to suppress viral titers (**Figure 9A, ii and iii, and Supplemental Figure 10, A and E**). No differences in viral mutation load were observed across ages (**Supplemental Figure 10F**), nor were differences in kinetics or magnitude of IFN gene responses, LDHB, or HMGB1 release detected (**Supplemental Figure 10, B and C**). These data are different than a smaller *in vitro* nasal epithelial study (57) but are consistent with age-dependent data from bronchial epithelia *in vitro* (29, 58) and *in vivo* nasal SARS-CoV-2 RNA measures in the respiratory system (14).

**Figure 9.**
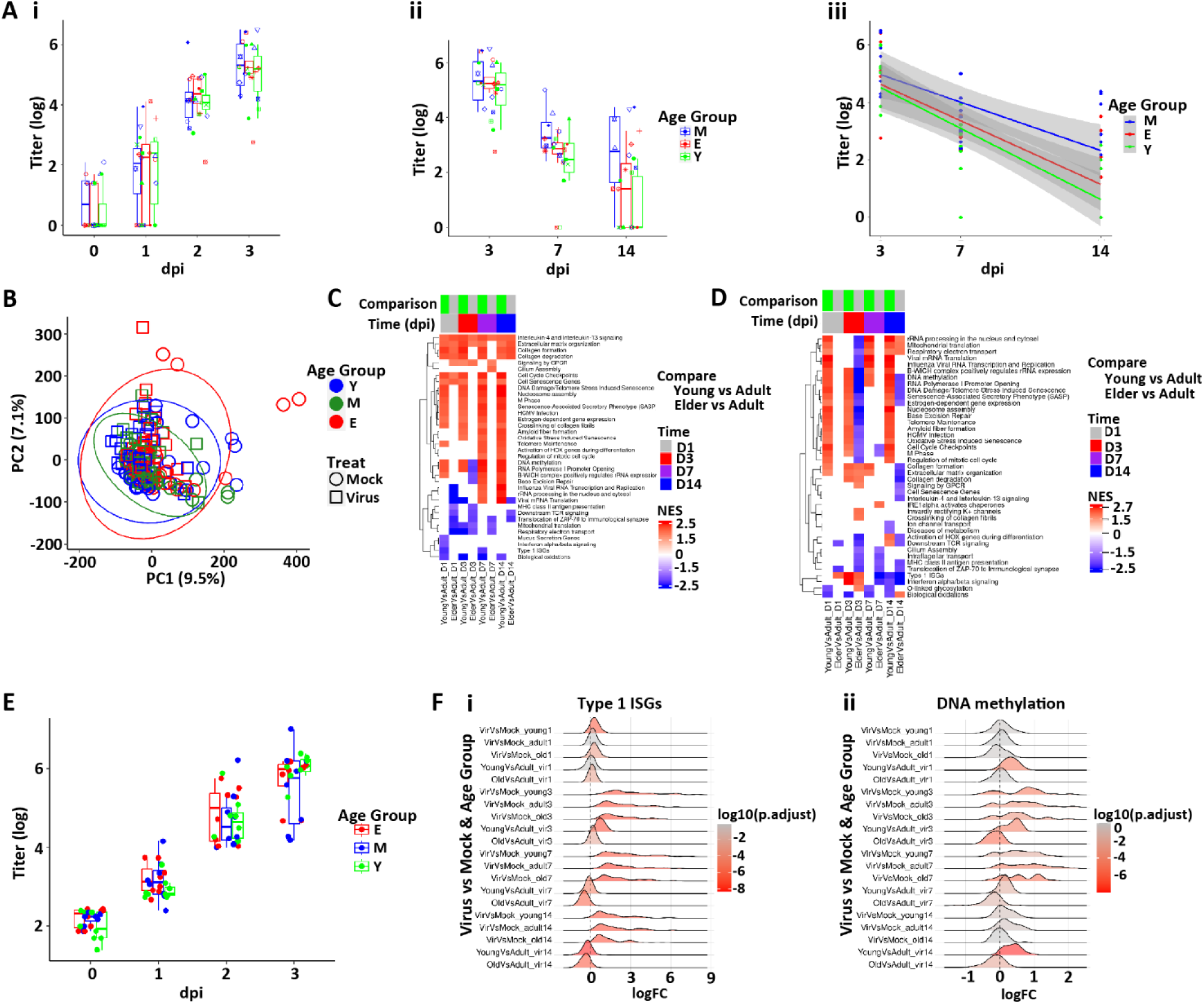
SARS-CoV-2 infectivity and HBE culture host responses as a function of age. **(A)** Box plots depicting mean titers in SARS-CoV-2 infected cultures separated into age groups. No significant differences were found among ages at any timepoint measured. **(Ai)** Titers 1-3 dpi; **(Aii)** titers 3-14 dpi. **(Aiii)** Slopes of viral titers from 3, 7, and 14 dpi as a function of age. M = mature; Y = young; E = elderly. Shaded areas represent the 95% confidence intervals. **(B)** PCA of global bulk RNAseq transcriptomics colored by age group. Ellipses are drawn at 95% level assuming normal distributions of samples per group. **(C)** Pathways selected from bulk RNAseq data by the Reactome + custom pathway annotation describing differentially regulated pathways in mock treated cultures between the age groups shown at the dpi indicated. **(D)** Selected differentially regulated pathways in virus-treated samples between the age groups shown at the indicated. For (C-D), NES=normalized enrichment score. **(E)** Replication Protocol 2 data for titer plotted by age. (**F)** Ridge plots representing **(Fi)** ISGs and **(Fii)** DNA methylation signatures between age groups and times dpi.

The PCA plot of the mock HBE RNAseq transcriptomic data showed no clear separation as a function of age, and surprisingly few differences in gene expression across age using our filtering criteria were found (**Figure 9D and Supplemental Figure 10 D**). We then sought age-dependent differences using pathway analyses in mock infected cultures across all timepoints (**Figure 9C**; see **Supplemental Table 2** for access to analyses). While each individual mock timepoints comparison produced pathways that survived statistical significance testing, the patterns were complex. A selected set of these pathways was chosen to depict major patterns (**Figure 9C, D**). A consistent upregulation was noted across time in the young vs. mature comparison in pathways that contained genes involved in cell cycle and DNA repair. Very few pathways were consistently downregulated in young vs. mature, a notable exception being biological oxidations. Overall, we conclude that the data supports the concept that cultures from donors of all ages are surprisingly similar at the gene expression level, consistent with the absence of age-dependent differences in infectivity in Protocol 1 and, indeed, the replication protocol (Protocol 2) (**Figure 9E**).

Evaluation of age-dependent transcriptional responses to SARS-CoV-2 infection with respect to epithelial damage and repair also yielded complex pathway responses with age across time post-infection (**Figure 9D)**. Again, few consistent differences across age were noted across the entire viral infection interval and ISG responses were similar across age groups (**Figure 9Fi**). The tendency for cell cycle genes to be higher in young observed in mock cultures was maintained after virus infection, except for 7 dpi where previous results show the highest cell cycling across all ages (**Figure 6B**). In contrast, some of these genes were significantly downregulated in elderly vs. mature after SARS-CoV-2 infection, indicating that the elderly may have reduced capacity to cycle/repair after injury than young and mature cultures. The young donor cultures also exhibited pathways indicating more robust DNA methylation (**Figure 9Fii**) with the elderly showed reduced responses in these general pathways compared to the mature (or young) group.

In sum, we conclude that the respiratory epithelia of all ages exhibit similar baseline gene expression properties and SARS-CoV-2 infectivity, but age-dependent differences in pathways associated with repair were manifest.

### J. Variables related to abnormal airway repair at 14 dpi in vitro

The contributions of peak infection intensity (titer, TPM, NCAP) and epithelial damage (LDHB) to a pathway-based index of failed epithelial repair (referred to as the repair index; see **Supplemental Methods**) were tested. Surprisingly, no relationships between the intensity of viral infection (**Figure 10A and Supplemental Figure 11, A and B**) and epithelial damage (**LDHB; Figure 10B**) and the 14 dpi repair index were detected. Similarly, no relationships were detected as a function of the sex of the donor (**Figure 10C**). In contrast, young age appeared to be associated with an improved repair index (**Figure 10D**).

**Figure 10.**
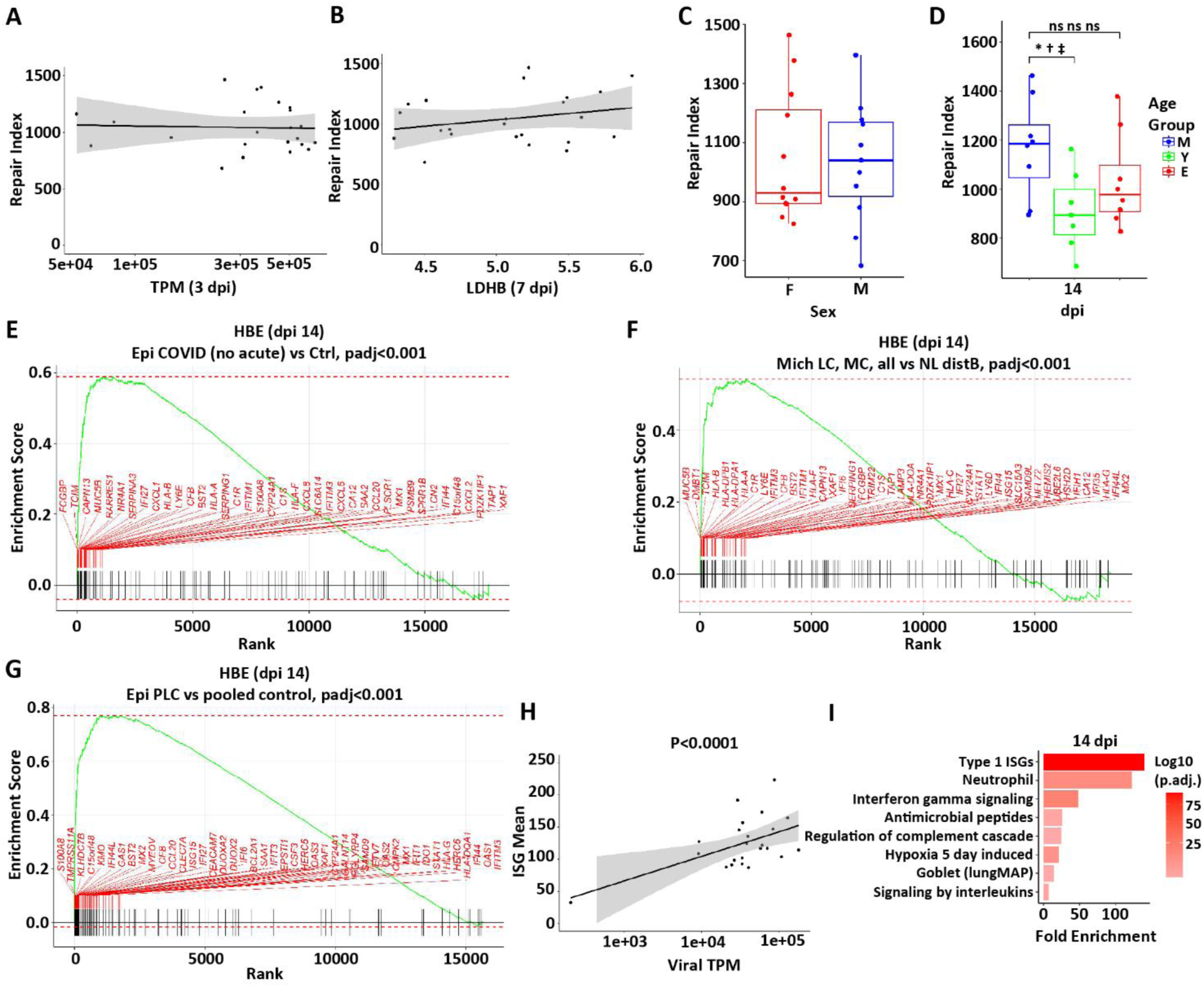
Relationship of failed repair to COVID-19 disease. A repair index score as a function of **(A)** viral gene expression (TPM) at 3 dpi, **(B)** peak LDHB protein in secretions at 7 dpi, **(C)** sex, and **(D)** age. All comparisons depict the repair index at 14 dpi. **(E-G)** Failed repair genes are enriched in GSEA analyses for **(E)** a reported autopsy (∼ 30 d post-SARS-CoV-2 symptom initiation) with early pulmonary fibrotic histologic changes (46); **(F)** GeoMX spatial transcriptomics studies of conducting airway epithelia of five subjects 6-12 mo post-initiation of SARS-CoV-2 symptoms who exhibited persistent CT abnormalities and required VATS biopsies, again with fibrotic changes evident on histology; and **(G)** transbronchoscopic epithelial biopsies of subjects exhibiting a long pulmonary COVID phenotype that manifests as chronic cough, sputum production, and chest congestion persistently for ∼ 18 mo post-SARS-CoV-2 infection vs. subjects who resolved symptoms post-SARS-CoV-2 (46). **(H)** Relationship between ISGs and viral TPM at 14 dpi (slope=0.68 SE 0.14, p<0.0001). **(I)** Selected pathways over-represented (enriched) in failed repair gene set. Shaded areas in A, B, and H represent the 95% confidence intervals.

### K. Repair processes reflect *in vivo* COVID

We next asked whether genes that reflected failed return to a normal state, *i.e.*, failed repair, at 14 dpi in HBE cultures exhibited relevance to persistent long-COVID pulmonary disease. Analyses of virus-infected vs. mock cultures identified 148 DEGs using an established threshold for significance (see **Supplemental Methods**). We utilized this 14 dpi gene signature of “failed repair” to query three datasets to address this question across two post-COVID lung disease phenotypes.

We first analyzed spatial transcriptomics data from two cohorts with a post-COVID pulmonary fibrosis phenotype. The first analysis focused on a reported autopsy study of 61 subjects ∼ 30 d post-SARS-CoV-2 symptom initiation with early pulmonary fibrotic histologic changes (46). GSEA analyses revealed significant enrichment for the 14 dpi repair genes (e.g., *FCGBP, MUC5B, SERPINA3, IFI27, CLXL1*) in persistently abnormal airway epithelia in this cohort (**Figure 10E**). The second analysis focused on GeoMX spatial transcriptomics studies of conducting airway epithelia of five subjects 6-12 mo post-initiation of SARS-CoV-2 symptoms who exhibited persistent CT abnormalities and required video-assisted transthoracic surgery (VATS) biopsies (**Supplemental Methods**). Histological evaluation of the biopsies revealed a pulmonary fibrotic phenotype. Notably, the GSEA analysis again revealed significant enrichment of the 14 dpi repair genes in diseased airway epithelial regions of the biopsied long-COVID lungs (**Figure 10F**). Thus, the altered repair processes identified in 14 dpi HBE persist in the airway regions of subjects with long-COVID pulmonary fibrotic phenotypes.

We next studied a long pulmonary COVID phenotype that manifests as chronic cough, sputum production, and chest congestion. Recently, transbronchoscopic epithelial biopsies of subjects exhibiting this phenotype persistently for ∼ 18 mo post-SARS-CoV-2 infection vs. subjects who resolved symptoms post-SARS-CoV-2 were reported (59). The 14 dpi HBE SARS-CoV-2 DEG signature was enriched in the symptomatic as compared to non-symptomatic/control post-SARS-CoV-2 infection groups, suggesting persistently abnormal repair contributes to the long-COVID bronchitic phenotype **(Figure 10G**). A pathway analysis was performed on the gene set representing failed repair. As predicted, the genes representing the ISG pathway were enriched and correlated with the presence of viral transcripts (**Figure 10H**). Other upregulated pathways in this gene set included neutrophil, complement, hypoxia, and interleukin signaling (**Figure 10I**).

## Discussion

The clinical outcomes of SARS-CoV-2 infections vary widely, reflecting a complex interplay between the virus and host defense and reparative properties. This study explored potential age- and sex-dependent contributions of respiratory epithelia to SARS-CoV-2 outcomes through 14 days post-infection. This study also offered the opportunity to investigate relationships between *in vitro* intermediate-term repair (14 d) with long-term epithelial properties in subjects with long-COVID respiratory syndromes.

No differences in the rates of acquisition or peak viral titers, TPM, viral proteins, or viral mutation load were consistently observed across cultures derived from children, mature adults, or the elderly, consistent with previous reports of bronchial epithelia (29, 58). Key cell biologic variables reported to govern infection rates also did not differ across age groups, including ACE2 expression, TMPRSS2 expression, culture ciliation, or kinetics/magnitude of IF responses, again consistent with previous reports (29, 30, 60). RNA bulk-seq analyses of paired mock HBE cultures prior to infection showed little differential gene expression in single genes across all donor ages in PCA plots and DEG analyses. However, pathway analyses revealed an upregulation in the young referenced to maturity of cell cycling and DNA repair pathways.

While all age groups exhibited similar epithelial morphology at d14, dominated by mucus metaplasia, the younger subject cultures exhibited more robust cell cycling, DNA repair, and oxidative stress pathway responses over the 3-14 dpi interval. Similar patterns of more rapid/efficient repair have been observed in young vs. elderly subject cultures after a variety of biologic and chemical insults (61), suggesting the young have in general more robust reparative capacities in response to injury. The relatively consistent gene expression patterns observed after virus across ages in these epithelial cultures suggest that immune cell function, not intrinsic epithelial function, likely account for the age dependence of viral titer and severity outcomes from SARS-CoV-2 infections (62–64). The childhood respiratory immune system has been reported to be more versatile/trained to respond to a SARS-CoV-2 infection and prevent spread to the lung with worsened clinical outcomes (13, 65–67). Conversely, a senescent immune system has been posited to account for the worsened outcomes in the elderly (62, 63).

Regarding sex, our aggregate studies revealed no differences in SARS-CoV-2 infection in female vs. male human bronchial epithelial cells, similar to a report of human nasal cultures in which no differences in SARS-CoV-2 titer were detected as a function of sex (68). Epidemiologic studies have clearly established that males, especially older males, exhibit more severe COVID-19 disease, but there is a paucity of large epidemiologic studies that have reported the differences in titer in males vs. females *in vivo* (9). The two available studies described mixed findings, with a small study of US Marine recruits reporting lower SARS-CoV-2 Ct values for males (9), whereas a larger Saudi Arabian study reported a significantly lower Ct value in females of all ages than males (62, 63).

The absence of a large effect of sex on SARS-CoV-2 epithelial infectivity is consistent with analyses of female vs. male in mock (non-infected) cultures. PCA or DEG analyses of bulk-RNAseq data revealed surprisingly few differences, limited to X and Y chromosomes, in baseline gene expression, including ACE2 and TMPRSS2, between male and females (69). Note, the Human Cell Atlas has released data describing multiple pulmonary epithelial cells as a function of sex (70). Like our study, no differences in gene expression were observed, suggesting human respiratory epithelia are not sex hormone responsive.

Combining the age and sex data into a larger dataset permitted insights into the airway epithelial cell biologic response to the acute infection and repair processes. Aspects of the acute 3 dpi response have been widely reported, including the selective infection of ciliated cells with apoptotic and likely necrotic cell death (4, 28, 71, 72). The cell signaling processes during this period are complex, with DAMP release, *e.g*., HMGB1 and ATP, but also recruitment of anti-inflammatory extra-cellular signals, *e.g*., MTA. Major cell biologic responses at this peak period also included suppression of mitochondrial and ribosomal gene expression. Coincident with these responses were striking increases in expression of genes associated with histone acetylation and inflammation. These responses reflect complex signaling cascades involving infected cells and communication *in vivo* with immune/inflammatory cells via IF, DAMP, and inflammatory mediators/cytokines.

Post-viral bacterial infections in the quasi-acute setting (wks) are common with respiratory viruses in general, and severe COVID-19 subjects have an ∼ 30% incidence of bacterial superinfections (73). Many of these bacterial infections involve infection of airway mucus plugs (46). Singanayagam et al., utilizing a rhinovirus challenge model in COPD subjects, demonstrated that post-viral bacterial infections reflected neutrophil destruction of host airway secretion antimicrobial substances and abnormal mucus production (74). Mucociliary clearance is a key defense mechanism combating airway infection and exhibits a complex interplay between ciliated cell numbers and mucus properties. Mucin secretion was upregulated, likely in part due to EGFR and IL1R1 signaling, in SARS-CoV-2-infected HBE cultures (46). However, mucins must be well hydrated to be transported effectively by MCC or the backup cough mechanism. The upregulation of the dominant ion channel mediating fluid secretion, *i.e*., CFTR, coupled to an absence of upregulation of the channel that mediates opposing fluid absorption, *i.e*., ENaC, generates a scenario whereby newly secreted mucus is sufficiently hydrated to mediate effective mucus clearance. However, the reduction post-SARS-CoV-2 infection in ciliated cell numbers, with a concomitant loss of cilial force and mucus hydration mechanosensing, can produce counterposing effects on MCC and cough clearance, particularly in small airways, that produce mucus plugging in COVID lungs (46).

Infection of mucus plugs may be promoted by the absence of activities of antimicrobial substances required for bacterial suppression on airway surfaces. In addition to neutrophil cleavage of these substances post-viral infection, SARS-CoV-2 infection downregulated in airway epithelia a complex of antimicrobial peptides, including CCSP, lactoferrin, and lysozyme over the 3-14 dpi interval (74, 75). These molecules are key for suppressing bacterial growth over intervals required for bacterial clearance by the MCC system, and the limitation of secreted antimicrobial substances in the context of hampered MCC provides favorable conditions for secondary airway bacterial infections (76).

Our study design allowed for detailed analyses of epithelial repair over 14 dpi in response to SARS-CoV-2 infection. Airway epithelia emerged from peak infection into a phase at 7 dpi of persistent but waning SARS-CoV-2 infection and repair/disrepair. By 14 dpi, airway epithelia repaired via remodeling pathways that produced an abnormally mucus metaplastic, less ciliated epithelium. This result likely is mediated in part by an increase in basal cell activation/cycling and a skewing toward a mucus metaplastic phenotype due in part to increases in EGF ligand production, *e.g*., amphiregulin, and EGF-R signaling (48, 49). Our data describing the arrest of ciliated cell maturation at the deuterosomal stage are consistent with this notion (40, 77). The major variable identified in our study that associated with repair at 14 dpi was age, with young repairing better, whereas no differences based on infection intensity or sex were detected.

Presumably, in most people post-SARS-CoV-2 infection, the epithelium continues to be repaired over weeks *in vivo* with resolution of symptoms. However, a fraction of people post-SARS-CoV-2 infection manifest “long pulmonary COVID” syndromes. Although multiple phenotypes are reported associated with long pulmonary COVID, chronic bronchitic and pulmonary fibrotic phenotypes are amongst the most common (59, 78–80). One mechanism that may contribute to long-COVID pulmonary disease is the failure of infected pulmonary epithelia to repair. This mechanism almost certainly contributes to the failure of infected alveolar AT2 cells to replenish both AT2 and AT1 cells in long-COVID subjects with a resultant chronic fibrotic phenotype (81–84).

Our data demonstrating the persistence of the HBE 14 dpi DEG signature in chronically diseased airway epithelia in people with long-COVID fibrotic and bronchitic phenotypes suggest persistent abnormal airway epithelial repair contributes to the pathogenesis of the airway component of long-COVID pulmonary disease. Analyses of the pathways characterizing this repair DEG set identified persistent ISG responses that correlated with the persistence of viral transcripts in HBE cells at 14 dpi. It has been previously shown that chronically elevated ISG expression in response to poly-IC exposure is associated with airway epithelial metaplasia, similar to reports in alveolar epithelia (85, 86). One speculation that may describe persistent ISG expression and airway metaplasia in some long-COVID individuals is a persistent low level of non-infectious SARS-CoV-2 RNA expression in airway epithelial cells not cleared via immunologic clearance mechanisms.

Our study has limitations. First, our study employed cultures of well-differentiated airway epithelia. Whereas this design queries the intrinsic activity of the epithelium, it does not reprise the activity of an epithelium *in vivo* exposed to the full spectrum of immune and stromal cells, nor can it reflect the normal clearance of virus via immune cell activity. Thus, the culture approach may have accounted for failure to identify key genes related to age-dependent host antiviral defense *in vivo*. Second, our cultures were not exposed to media containing sex hormones that may have been appropriate for sex. Third, the experimental sampling intervals were not daily, making interpretation of rates of accumulation of virus/secreted proteins difficult. Normalization was used to mitigate this problem. Fourth, while relatively large for a study in cell culture, the study is expected to be underpowered to capture the entirety of the variability in the human population.

In summary, our studies demonstrate that SARS-CoV-2 infection of airway epithelia leaves the airway epithelium at 14 dpi in a repairing but complex state characterized by impaired mucus transport, host defense protein secretion, and hyperinflammation. Young age, but neither sex nor the peak intensity of the SARS-CoV-2 infection, modulates the post-viral repair state, consistent with improved outcomes of the young (17, 26, 67, 87). The incomplete/abnormal repair state is predicted to render the airway epithelium over a quasi-acute (days) post-infection interval vulnerable to bacterial superinfections. The persistence of a failed repair state over the longer term (months) may contribute to post-COVID pulmonary syndromes, including an airways component of fibrotic sequelae and a bronchitic phenotype of cough and sputum production. A persistent airway epithelial ISG response, possibly in response to poorly cleared viral RNA, may be a driver of aspects of these long-COVID syndromes.

## Methods Section

Our study utilized both male and female human airway epithelial cultures at equal proportion, and sex differences were specifically queried.

### A. Experimental Design for *In Vitro* Studies

The experimental design is outlined in **Supplemental Figure 1**, with the technical details of each individual assay provided in the **Supplement Material**. In broad terms, air–liquid interface (ALI), well-differentiated HBE cultures were infected with SARS-CoV-2 and collected for analyses as indicated (88). Cells were obtained under the protocol #03-1396 approved by the University of North Carolina at Chapel Hill Biomedical Institutional Review Board in the Marsico Lung Institute Tissue Procurement and Cell Culture Core. Three age groups were studied: Young (age 0-8); Mature (age 21-29): Elderly (> 69). A replication protocol studied the same donors in a single batch over a 3 dpi interval (**Supplemental Figure 1B**). Sample specifics are provided in **Supplemental Table 1.** Key assays conducted consisted of titer, bulk RNAseq, and proteomics from apical secretions. Key analyses included determination of differential gene and protein expression (including pathway analyses), development of heatmaps, and an index to describe disease repair. The index was tested against candidate variables hypothesized to govern repair status, including infection intensity, sex, and donor age. Differences of means of outcome measurements between experimental groups were tested with linear models, or in the case of repeated measures design, with linear mixed-effect models. Analyses methods are provided in detail in **Supplemental Material,** with details provided in figure legends as appropriate. Full analyses of the results are provided in **Supplemental Files 1-24** with a description of those files provided in **Supplemental Table 2**.

## Data Availability

Underlying bulk RNAsequencing data for this study have been submitted to the Gene Expression Omnibus (GEO) repository and are publicly available at https://www.ncbi.nlm.nih.gov/geo/query/acc.cgi?acc=GSE299837. All other data associated with this manuscript are included as Supplementary Files (see **Supplemental Table 2** for details). The data points utilized for each plot are also available in the Supporting Data Values file accompanying this publication.

## Acknowledgments

The authors wish to thank Eric Roe at the Marsico Lung Institute for expert editorial and administrative assistance in the preparation of the manuscript. We also acknowledge the University of North Carolina Pathology Services Core for providing histological samples, especially the director of this core, Dr. Gabriela De la Cruz and her staff.

