## Supplemental Table 1 for "Airway epithelial SARS-CoV-2 infectious and repair responses: relationships to age, sex, and post-COVID pulmonary syndromes"

**Supplementary Table 1.** Features of cell cultures used in this study.

| Donor No. | Donor ID | Batch** | Age (yr)$ | Sex | Ancestry |
|---|---|---|---|---|---|
| 1 | Y-58P | 1 | 4 mo | M | Caucasian |
| 2* | Y-15K | 1 | 10 mo | M | Caucasian |
| #3 | Y-37O | 1 | 2 | M | Caucasian |
| 4 | Y-48G | 1 | 5 | F | Caucasian |
| 5 | Y-49O | 1 | 8 | F | Hispanic |
| #6 | M-9N | 1 | 21 | M | African American |
| 7 | M-10H | 1 | 26 | M | Caucasian |
| #8 | M-25O | 1 | 26 | F | Caucasian |
| 9 | M-19O | 1 | 27 | M | Unknown |
| #10 | M-74K | 1 | 29 | M | Caucasian |
| 11 | E-5I | 1 | 69 | M | African American |
| 12 | E-10G | 1 | 70 | F | Caucasian |
| #13 | E-35K | 1 | 70 | M | Caucasian |
| 14 | E-26P | 1 | 73 | F | Caucasian |
| #15 | E-3N | 1 | 91 | M | Caucasian |
| 16 | Y-18I | 2 | 2 mo | F | Caucasian |
| 17 | Y-46M | 2 | 4 mo | F | Caucasian |
| 18 | Y-83K | 2 | 10 mo | M | African American |
| 19 | M-35P | 2 | 21 | F | Caucasian |
| 20 | M-51K | 2 | 21 | F | Hispanic |
| 21 | M-21M | 2 | 23 | F | Caucasian |
| 22 | E-41T | 2 | 69 | F | Unknown |
| 23 | E-30T | 2 | 70 | M | Unknown |
| 24 | E-11G | 2 | 78 | F | Caucasian |

*This sample was excluded from the Protocol 1 dataset due to experimental/technical errors during viral infection.
**This batch number refers to Protocol 1 only; Protocol 2 was conducted in one batch.
NOTE: in addition to the above footnote, the following two samples were excluded from the RNAsequencing analyses due to failure to pass QC testing: Donor #1, 1 dpi mock; and Donor #11, 14 dpi mock.
#For the study of metabolites, these donors were used.
$mo=months for donors < 1 year old.
